# Dysfunctional temporal stages of eye-gaze perception in adults with ADHD: a high-density EEG study

**DOI:** 10.1101/2021.10.14.464365

**Authors:** Cheyenne Mauriello, Eleonore Pham, Samika Kumar, Camille Piguet, Marie-Pierre Deiber, Jean-Michel Aubry, Alexandre Dayer, Christoph M. Michel, Nader Perroud, Cristina Berchio

**Affiliations:** Department of Psychiatry, Faculty of Medicine, University of Geneva, Geneva, Switzerland; Department of Psychology, University of Cambridge, Cambridge, United Kingdom; Department of Basic Neurosciences, Faculty of Medicine, University of Geneva, Geneva, Switzerland; Department of Mental Health and Psychiatry, Service of Psychiatric Specialties, Mood disorders unit, University Hospitals of Geneva, Switzerland

**Keywords:** ADHD, ERP, face perception, eye-gaze

## Abstract

ADHD have been associated with social cognitive impairments across the lifespan, but no studies have specifically addressed the presence of abnormalities in eye-gaze processing in the adult brain.

This study investigated the neural basis of eye-gaze perception in adults with ADHD using event-related potentials (ERP). Twenty-three ADHD and 23 controls performed a delayed face-matching task with neutral faces that had either direct or averted gaze. ERPs were classified using microstate analyses.

ADHD and controls displayed similar P100 and N170 microstates. ADHD was associated with cluster abnormalities in the attention-sensitive P200 to direct gaze, and in the N250 related to facial recognition. For direct gaze, source localization revealed reduced activity in ADHD for the P200 in the left/midline cerebellum, as well as in a cingulate-occipital network at the N250. These results suggest brain impairments involving eye-gaze decoding in adults with ADHD, suggesting that neural deficits persist across the lifespan.

## 1. Introduction

Individuals who have Attention-Deficit/Hyperactivity Disorder (ADHD) struggle with inattention, hyperactivity and impulsivity (APA, 2013). ADHD is also associated with a range of social and interpersonal problems across the lifespan (Canu, Tabor, Michael, Bazzini, & Elmore, 2014; Gardner & Gerdes, 2015; Ronk, Hund, & Landau, 2011). Hyperactive and impulsive behaviors, as well as inattention, may cause problematic social functioning, including poor interactions or inappropriate social behaviors (Guntuku, Ramsay, Merchant, & Ungar, 2019; Michielsen et al., 2015).

Although the core symptoms of ADHD may contribute to the emergence of hindered social interactions, a range of social cognitive impairments have been also associated with ADHD (Uekermann et al., 2010). ADHD individuals show deficits related to understanding social signals, such as reading states of mind (Mary et al., 2016; Tatar & Cansız, 2020), and recognizing facial expressions (Schönenberg, Schneidt, Wiedemann, & Jusyte, 2019). Eye-gaze signals are critical to decipher social and affective signals (Adams Jr & Kleck, 2005; Becchio, Bertone, & Castiello, 2008; Capellini, Riva, Ricciardelli, & Sacchi, 2019). Eye-gaze signals create social-affective coordination by stating approach or avoidance tendencies (Adams Jr & Kleck, 2005; Hietanen, 2018), as well as informing about the focus of attention (Pierno et al., 2006). Unsurprisingly, children and adolescents with ADHD have difficulties processing eye-gaze signals (Airdrie, Langley, Thapar, & van Goozen, 2018; Guo et al., 2019; Marotta et al., 2014). It is unknown whether these early abnormalities persist into adulthood since, to our knowledge, no study on eye-gaze processing in adult ADHD patients has been published.

Event-related potentials (ERPs) enable the study of discrete cognitive processes related to face and gaze perception with high temporal resolution. ERP components are sensorial/cognitive evoked responses, temporally and spatially defined by latency and scalp field configuration. The first visual evoked response to faces occurs with a posterior positivity at around 100 ms (P100)(Herrmann, Ehlis, Ellgring, & Fallgatter, 2005; Seki et al., 1996), and is already modulated by attention (Eimer, Holmes, & McGlone, 2003). At approximately 170 ms after face presentation, a latero-posterior negative component corresponds to the encoding of facial features in the human brain (N170) (Bentin, Allison, Puce, Perez, & McCarthy, 1996; Gao, Conte, Richards, Xie, & Hanayik, 2019; Hinojosa, Mercado, & Carretié, 2015). Middle latency stages (150-250 ms) reflect higher-order cognitive processes of face decoding (Olofsson, Nordin, Sequeira, & Polich, 2008), such as selective attention and facial identity recognition (P200/N250 (Lijffijt et al., 2009; Singhal, Doerfling, & Fowler, 2002)). ERPs appearing at later latencies (250-400 ms) usually reflect emotional stimuli processing (Bublatzky, Gerdes, White, Riemer, & Alpers, 2014; Stolz, Endres, & Mueller, 2019). By classifying ERP map configurations into distinct clusters, ERP microstate analyses describe dynamically varying scalp potential patterns and allow us to explore potential dysfunctional large-scale brain networks between groups (Michel & Murray, 2012). These networks can be further investigated using electrical neuroimaging, permitting the reconstruction of cerebral sources with high temporal resolution (Michel & Brunet, 2019).

Only a few studies using classical procedures of analysis have investigated face processing in adults with ADHD. In adult ADHD compared to controls, Raz and Dan (Raz & Dan, 2015) reported a reduced P300 amplitude only in response to face targets, while Ibáñez and co-workers (Ibáñez et al., 2011) found reduced N170 responses to emotional faces. Other authors have documented a reduced late positivity to negative emotions during response inhibition in ADHD (Köchel, Leutgeb, & Schienle, 2012). To our knowledge, only studies on children with ADHD have investigated the neuro-biological mechanisms of eye-gaze perception (Groom et al., 2017; Guo et al., 2020; Guo et al., 2019; Tye et al., 2013). Tye and co-workers (Tye et al., 2013) provided evidence of a reduced effect of eye and face inversion on the P100, suggesting that impairments in eye-gaze processing might manifest as an initial visual/ vigilance bias toward social signals. Guo and co-workers showed that children with ADHD have an inverse posterior alpha pattern to eye-gaze signals, predictive of inattention (Guo et al., 2019).

The aim of the current study was to investigate the neuro-psychological basis of eye-gaze perception in adults with ADHD. To measure early stages of face processing, we used high-density ERP and a validated delayed face-matching task (Berchio et al., 2016), in which faces with direct and averted gaze were presented.

On the basis of previous works suggesting that children and adolescents with ADHD exhibit atypical processing of eye-gaze signals (Airdrie et al., 2018; Guo et al., 2019; Marotta et al., 2014), we hypothesized that similar impairments would be observed in adults with ADHD. Specifically, we hypothesized that ADHD patients would show reduced visual orienting to social signals associated with P100 abnormality, altered face encoding responses revealed by N170 alteration, and atypical attentional differentiation to direct vs. averted gaze indexed by middle processing stages’ components. As there are no neuroimaging studies that have previously investigated the effect of gaze direction on face perception in ADHD, no specific predictions were made for this particular aspect. Although some behavioral studies failed to find differences between adults with ADHD and healthy controls in face recognition (Borhani & Nejati, 2018; Ibáñez et al., 2011; Noordermeer et al., 2020), we expected to observe shorter reaction times and more mistakes to face recognition in individuals with ADHD, mainly due to symptoms of impulsivity and inattention.

## 2. Methods

### 2.1 Participants

Twenty-three individuals with ADHD and 23 healthy control participants were included in the study. ADHD patients were outpatients from the Geneva University Hospital (HUG), followed in the Emotional Dysregulation Unit (TRE) for adult ADHD program from the Psychiatry Department. Psychiatric diagnoses in all subjects were established with the Diagnostic Interview for Genetic Studies (DIGS) as assessed by trained psychologists (EP, ALK).

During a clinical visit by a psychiatrist or psychologist trained in the evaluation of ADHD, patients were assessed using the ADHD Child Evaluation for Adults (ACE+), a semi-structured interview developed to support healthcare practitioners in the assessment and diagnosis of adults with ADHD (freely available at: https://www.psychology-services.uk.com/adhd.htm), and the French version of the Diagnostic Interview for Genetic Studies (DIGS, mood disorder parts only (Preisig et al., 1999).

Control subjects were matched for age, gender, and laterality, and they were recruited through advertisements placed at the University of Geneva and on classified websites. Exclusion criteria for all participants were any history of head injury or mental retardation, as well as current alcohol or drug abuse. Controls were excluded if they had a history of psychiatric or neurological disease, as assessed during an interview.

Sixteen patients were taking psychostimulants (methylphenidate), but all of them were requested to discontinue medications for the 24 hours prior to EEG recording, as previously described (Deiber et al., 2020).

Quantification of current symptoms was performed using the standardized and validated Adult ADHD Self-Report Scale for adults (ASRS-v1.1 (Kessler et al., 2005)), which consists of two subscales evaluating symptoms of attention deficit disorder, and hyperactivity/impulsivity. State and trait anxiety were assessed using State–Trait Anxiety Inventory (STAI; (Spielberger, 1983)). To exclude potential biases due to short-term memory deficits in ADHD, two working memory indexes were assessed: digit span and arithmetic (Wechsler, 2008).

All participants gave written informed consent prior to assessment. The research was conducted according to the principles of the Declaration of Helsinki and approved by the University of Geneva research ethics committee (CER 13-081). Participants received vouchers from a general store in exchange for their participation in the study.

### 2.2 Experimental paradigm

To assess neural mechanisms of spontaneous eye-gaze processing, participants performed a validated delayed face-matching task (Berchio et al., 2016). Stimuli were neutral faces (female: 50%) with direct or averted gaze. Each unique face had a single eye-gaze direction for the entire task. The task was to identify as quickly and accurately as possible whether a presented face was the same as the one shown two faces before. This paradigm has been developed to induce spontaneous mechanisms of eye-gaze recognition (Berchio et al., 2016). Participants used their right hand to press a down-arrow key if the stimuli matched and an up-arrow key if they did not match.

Faces were displayed on the centre of the screen for one second, with a fixed inter-stimulus interval of two seconds (during which a white fixation cross appeared). We considered a “target face” (i.e., the face stimulus presented with direct or averted gaze) that was identical to the one presented two faces before as a “match-trial” (or “match face”). A mismatch with the target face was called a “mismatch-trial”. Match-trials and mismatch-trials were presented with a 40:60 ratio. Three blocks of 120 trials were administered, each block consisting of 60 direct-gaze-face trials and 60 averted-gaze-face trials presented pseudo-randomly. Each block lasted five minutes, and the total task duration was 15 minutes. Before the task, participants performed a short training block of 10 trials. During the task, participants were seated at a table with the head resting on a chin rest, at a viewing distance of 60 cm from the screen.

### 2.3 EEG data acquisition and pre-processing

EEG data were acquired at 1000 Hz using a 256-channel system (EGI, Philips Electrical Geodesics, Inc.), in a lighted Faraday-cage room. Electrode impedance was kept below 30 kΩ, and data were acquired with the vertex (Cz) as reference electrode.

ERP data time-locked to face presentation onset, for correct trials only, were averaged separately for the two target conditions and the two groups. Data were band-pass filtered between 0.4 and 40 Hz, and a 50-Hz notch filter was applied. The original montage was reduced from 256 to 204 channels to exclude face/neck channels located at the periphery (see (Berchio, Rodrigues, Strasser, Michel, & Sandi, 2019)). Epochs for ERP analysis began 100 ms before fixation cross and ended 600 ms after stimulus onset. Epochs contaminated by artifacts (muscle, eye-blink/movements) were excluded by visual inspection. Epochs were re-referenced to the average, baseline-corrected against the mean voltage of the 100-ms pre-stimulus period, and down-sampled to 250 Hz. Data pre-processing was performed using CARTOOL Software programmed by Denis Brunet (Brunet, Murray, Michel, & neuroscience, 2011).

There were no group differences in the number of trials accepted (ADHD: direct gaze: *M*= 45.17, *SD*=7.67, averted gaze: *M*=47.52, *SD*=8.02; Controls: direct gaze: *M*=46.91, *SD*=6.44, averted gaze: *M*=49.52, *SD*=6.63; *p >* 0.36. two-tailed –t-test).

### 2.4 Behavioral analyses

Behavioral response data were analyzed in the framework of the signal detection theory, as measured by the discriminability index (*d’*), and the criterion (*C*) (Macmillan, 1993; McNicol, 2005). These parameters were assessed as an index of the participant’s ability to discriminate eye-gaze shifts. *D’* enables an estimate of the capability to differentiate the signal from the noise, and it was calculated as: *d’* = Z(false alarms + misses) – Z [(hit rates + correct rejections) + (false alarm + misses)]. *C* reflects the strategy of response of being more willing to say yes rather than no, and it was calculated as: *C* = (–1/2) · [z (hit rates) + z (false alarms)]. A greater tendency to report signal is therefore indicated by C values less than zero, a reduced tendency by values greater than zero. *D’* and *C* for the delayed face-matching task were calculated. Behavioral performance was also assessed using Pashler’s K as an index of working memory capacity, which was calculated as: K = N (hit rates – false alarms) / (1 – false alarms) where N is the number of items to be stored (Rouder, Morey, & Morey, 2011).

Median reaction times (RTs) of correct answers (hit rates) were also calculated. For each behavioral measure (d’, C, K, RT), a repeated-measures ANOVA was conducted for comparisons between groups, with Gaze as within-subject factor (‘Direct gaze’ vs. ‘Averted gaze’) and Group as between-subject factor (‘ADHD’ vs. ‘Controls’). To clarify main effects and interactions, post hoc analysis used paired t-tests with *p* < .05 as significance threshold after Bonferroni correction for multiple comparisons.

### 2.5 ERP microstate analysis

To assess differences between groups, ERPs were analyzed using microstate analyses (Michel & Murray, 2012; Murray, Brunet, & Michel, 2008). This cluster-based approach, which is a reference-free method, is used to classify ERP components into stable scalp configurations. The primary assumption behind this approach is that when a map changes its spatial distribution, the active brain sources also change (Lehmann, 1987; Vaughan, 1982). Therefore, it can also be used to differentiate groups.

The microstate analysis was performed in three steps ((Pascual-Marqui, Michel, & Lehmann, 1995); for more details see also (Murray, De Lucia, Brunet, & Michel, 2009)). First, the four grand averages (two conditions, two groups) were submitted to a k-means clustering, a non-hierarchical procedure allowing to make hypotheses about potential differences between groups. To exclude small maps that are physiologically implausible, a rejection criterion of 20 ms was applied (i.e., minimum map duration). The optimal number of maps was determined by the meta-criterion implemented in Cartool, which defines the optimal number as the median across all applied criteria.

Second, to test the k-means assumptions, the maps that were identified by the clustering procedure were back-fitted to the individual ERP data. This procedure consists of re-assigning the maps with the highest correlation to the data, resulting in a classification of the data at each point in time with a given map.

This fitting procedure allows to extract mean correlation, global explained variance (GEV), time coverage (time frames), and mean duration (ms) of each map and individual subject. These variables were compared to assess differences between groups. Before choosing appropriate tests, the equivalence of variances between groups was evaluated. To correct for the multiple comparisons, as a certain degree of associations between variables was expected, we first estimated the number of independent tests (M) and adjusted the test criteria to M with the *Sidak* correction, and then the null hypothesis was rejected for any test where the *p*-values were higher than 0.009 (Li & Ji, 2005).

#### Brain source analyses

To assess differences between groups on brain activation networks, we applied a linear distributed inverse solution model (LORETA,(Pascual-Marqui, Esslen, Kochi, & Lehmann, 2002)). The inverse solution was estimated for each individual’s average ERP on a Locally Spherical Model with Anatomical Constraints [LSMAC model (Brunet et al., 2011)], on 5018 voxels located on the grey matter of the average brain template of the Montreal Neurological Institute (http://www.bic.mni.mcgill.ca/brainweb). A spatial filter was applied to reduce noise in the data; following the computation of the inverse solution, data were transformed by a modified z-score normalization implemented in Cartool (3.80) (for technical details see: (Michel & Brunet, 2019)).

To minimize type I error, group-level network analyses were performed using a randomization test implemented in Cartool, on regions of interest (ROIs) defined by the Automated Anatomical Atlas (AAL, (Tzourio-Mazoyer et al., 2002), 116 ROIs), with a *p* value less than .01, on an average temporal window of interest. To highlight the decrease/increase in activity, *t*-values were reported if the associated *p* values were significant. Time-windows for brain-source analyses were determined based on evidence of group differences in the ERP microstate analysis.

## Results

### Demographic and clinical variables

No differences regarding age [*F*(1, 45) = 0.560, *p* = .46], gender [χ2(1) = 0.000, = 1], or handedness [χ2(1) = 0.000, = 1] were observed between groups.

ADHD participants showed significantly higher scores on the ASRS-Inattention subscale [Mann–Whitney *U*= 30, *Z*=-5.16, *p*<.001; d_Cohen_=2.33], and Hyperactivity/impulsivity subscale [Mann–Whitney *U*= 61.5, *Z*=-4.46, *p*<.001; d_Cohen_=1.75] than controls. Differences between groups for STAI-state [Mann–Whitney *U*= 105, *Z*=-3.51, *p*<.001; d_Cohen_=1.20] and STAI-trait [Mann–Whitney *U*= 75, *Z*=-4.16, *p*<.001; d_Cohen_=1.55] were also observed, indicating higher levels of state- and trait-anxiety in the ADHD group. No differences were observed between groups for digit span [*F*(1, 45) = 3.11, *p* = .08] or arithmetic [*F*(1, 45) = 1.16, *p* = .28]. Demographic and diagnostic information are summarized in Table 1.

**Table 1.** Demographic and clinical variables.

|  | **ADHD** |  | **Controls** |  |  |
| --- | --- | --- | --- | --- | --- |
|  | (n=23) |  | (n=23) |  |  |
|  | *Mean* | *SD* | *Mean* | *SD* | *P* |
| **Gender (n):** |  |  |  |  | n.s. |
| Male | 13 |  | 13 |  |  |
| Female | 10 |  | 10 |  |  |
| **Laterality (n):** |  |  |  |  | n.s. |
| Right-handed | 22 |  | 22 |  |  |
| Left-handed | 1 |  | 1 |  |  |
| **Age (years):** | 24.2 | 3.7 | 23.3 | 3.8 | n.s. |
| **Adult ADHD Self-Report Scale:** |  |  |  |  |  |
| Attention deficits | 24.6 | 5.3 | 11.6 | 4.9 | $p<.001$ |
| Hyperactivity disorder | 18.4 | 6.3 | 8.9 | 5.3 | $p<.001$ |
| **State-Trait Anxiety Inventory:** |  |  |  |  |  |
| STAI-state | 35.2 | 11.5 | 26.1 | 4 | $p<.001$ |
| STAI-trait | 45.9 | 9 | 34.2 | 7 | $p<.001$ |
| **WAIS-IV:** |  |  |  |  |  |
| Digit span | 10.34 | 1.7 | 11.28 | 1.8 | n.s. |
| Arithmetic | 13.78 | 2.6 | 14.72 | 3.2 | n.s. |

**Table 2.** Behavioral data.

|  | **ADHD** |  | **Controls** |  |
| --- | --- | --- | --- | --- |
|  | *Mean* | *SD* | *Mean* | *SD* |
| **d':** |  |  |  |  |
| Direct gaze | 1.999 | 0.467 | 2.033 | 0.416 |
| Averted gaze | 2.411 | 0.647 | 2.411 | 0.520 |
| **C:** |  |  |  |  |
| Direct gaze | -0.032 | 0.710 | -0.047 | 0.681 |
| Averted gaze | -0.025 | 0.648 | -0.011 | 0.594 |
| **K:** |  |  |  |  |
| Direct gaze | 2.21 | 0.450 | 2.41 | 0.340 |
| Averted gaze | 2.40 | 0.451 | 2.61 | 0.351 |
| **Reaction times (ms):** |  |  |  |  |
| Direct gaze | 678 | 95.8 | 711 | 86.8 |
| Averted gaze | 685 | 100.4 | 697 | 78.8 |

### Behavioral results

*D’* was significantly modulated by Gaze [F(1, 44) = 41.38, *p* < .001; η^2^=.485]. However, neither the main effect of Group nor the interaction between Group and Gaze were significant (all *p*_*s*_ > .78). Bonferroni-adjusted paired t-tests indicated that participants showed augmented sensitivity (*d*’) in recognizing faces with averted gaze than faces with direct gaze (*p* < .001). There were no significant main effects or interaction of Group and Gaze for the *C* criterion (*p*_*s*_*>* .82). We also computed K values to estimate groups’ memory capacity for each condition. In both groups, K values were significantly greater for faces with averted gaze than for faces with direct gaze [F(1, 44) = 41.38, *p* < .001; η^2^=.425] Bonferroni-adjusted paired t-test *p* < .001]. There were no significant main effects or interactions of Group and Gaze for the K values (*p*_*s*_*>* .08).

There was a marginally significant interaction of Gaze x Group [F(2, 44) = 3.630, *p* = .063; η^2^=.011] on the RTs. Data indicated that ADHD patients have a tendency to show faster RTs to faces with direct gaze when compared to controls.

### Face-evoked responses and microstate analyses

Based on visual inspection of the grand average ERPs (see Figure 2), four main evoked responses were recognizable for both conditions and groups: P100, N170, P200, and an enhanced negative potential in posterior electrodes approximatively 250 ms after stimulus onset (labelled N250).

**Figure 1.**
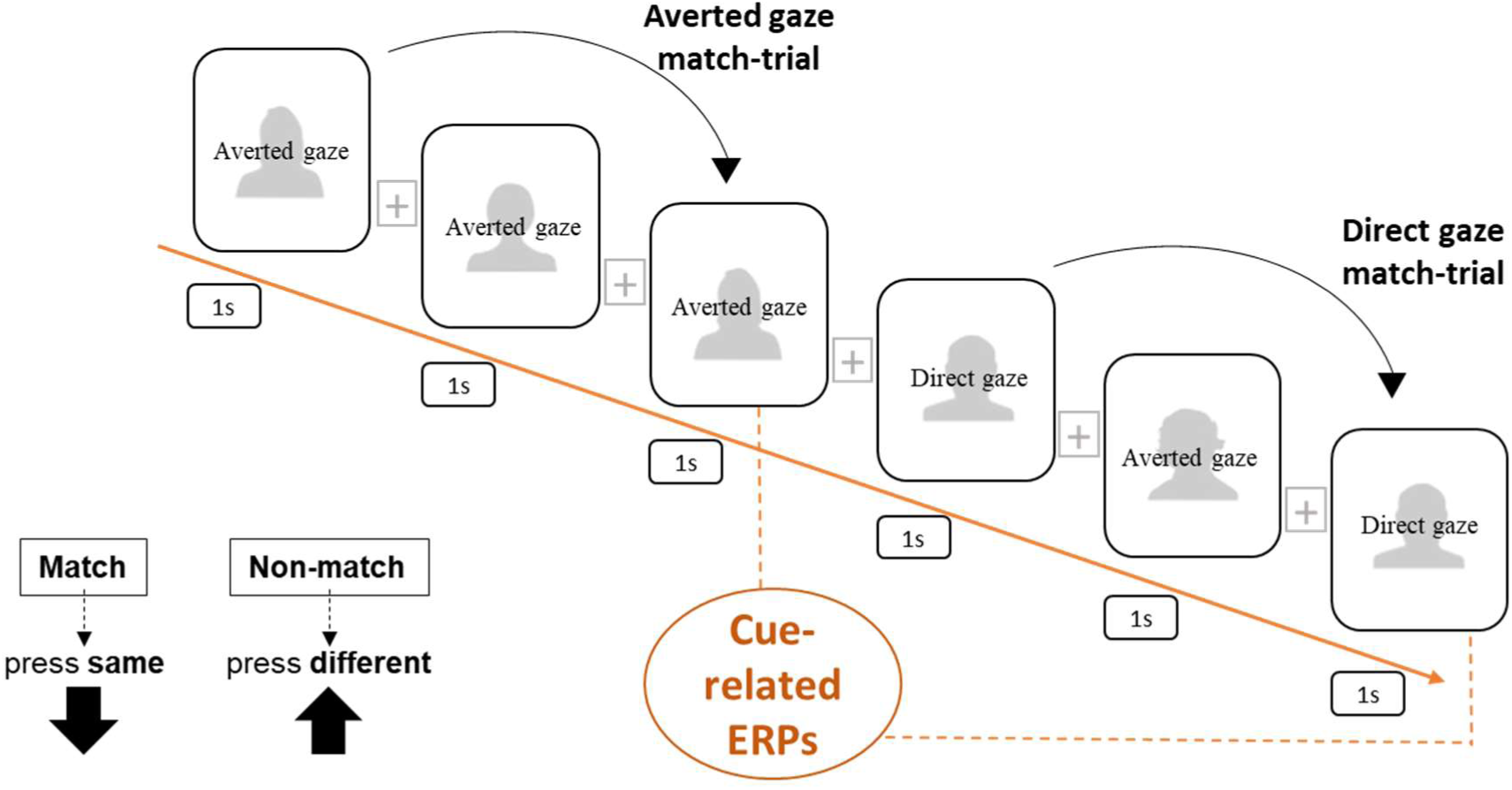
Schematic representation of the 2-back condition. Each stimulus face is presented for 1 s, followed by a grey display with white fixation cross for 2 s. Participants were asked to judge if each face was the same as the one appearing two trials before

**Figure 2.**
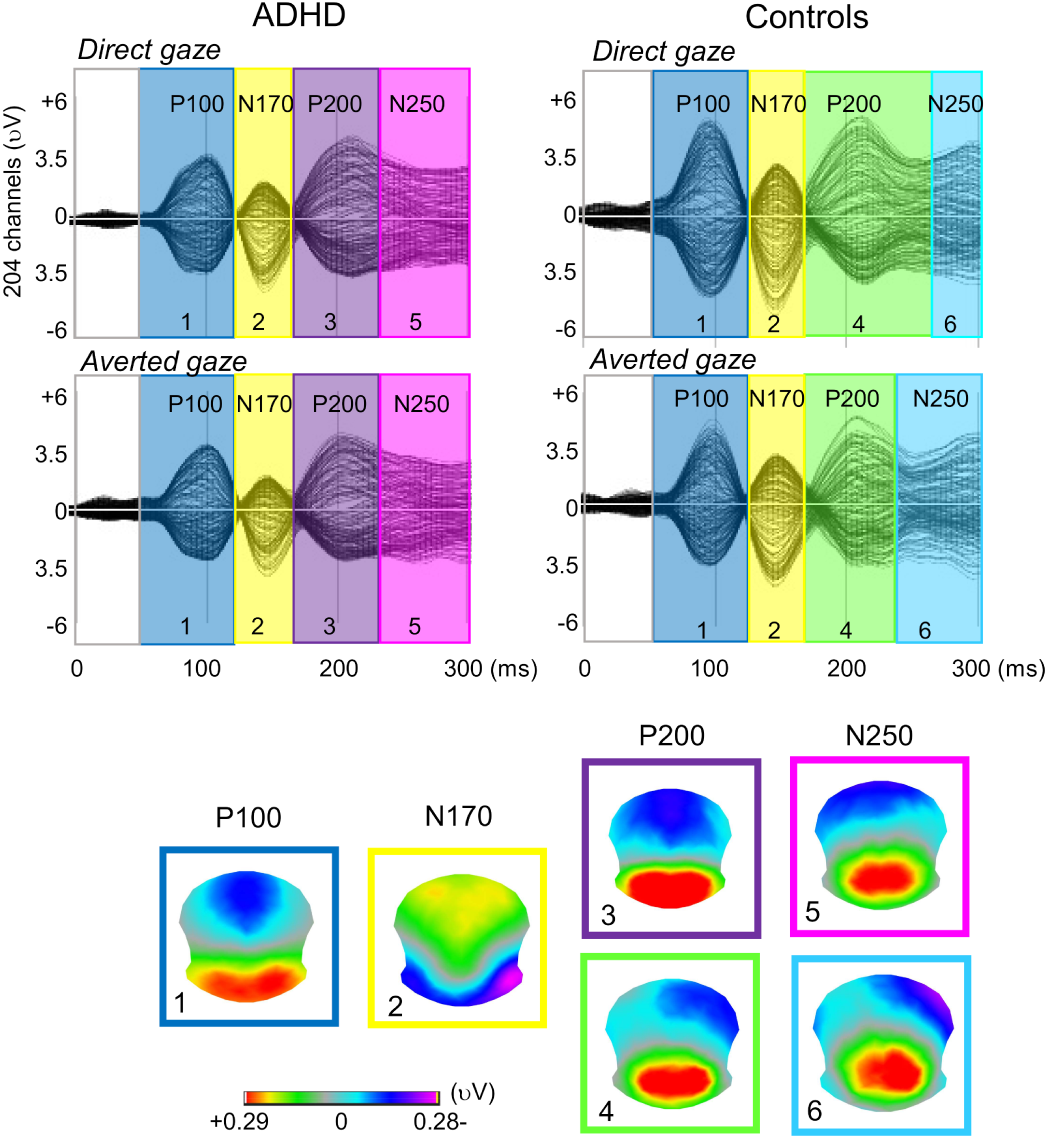
Evoked Responses for direct and averted gaze (grand averages, butterfly montage, 204 electrodes), for the ADHD patients and the controls. Four main components are elicited during the 300 ms post-stimulus: the visual P100, the face-related N170, the P200 that provides an index of attentional resources to eye-gaze cues, and the N250 which has been associated with facial identity recognition. Different background color stands for different microstate assignment. Map templates of each given microstate are shown in the lower part of the figure. To test whether group differences were statistically meaningful, the four maps that were identified as different between groups (3, 4, 5, 6) were fitted back to the individual ERP data at the corresponding time window of these components (172-300 ms). Since the majority of these variables were not normally distributed, non-parametric statistics (Mann–Whitney *U* test) were used for analyses. For all subjects and experimental conditions, microstate parameters are plotted in Figure 3.

Since the four main evoked responses were identified within the first 300 ms, k-means on grand averages was performed from 0 to 300 ms post-stimulus onset. This analysis revealed seven different classes of microstates: one for the baseline (map 0, 0-72 ms), one for the P100 (map 1; 72-124 ms), one for the N170 (map 2;124-168 ms), two for the P200 (map 3, 168-236ms; map 4, direct gaze: 168-276 ms, averted gaze: 168-232 ms), and two for the N250 (map 5, 236-300 ms; map 6, direct gaze: 276-300 ms, averted gaze: 232-300 ms). This first cluster analysis on the grand averages indicated potential differences between groups at both the P200 and the N250, with maps 3 and 5 identified for ADHD, and maps 4 and 6 for controls (see Figure 2).

For the P200 (maps 3 and 4), group differences were confirmed only for map 4 in the direct gaze condition. ADHD patients showed significantly lower map 4 values than controls for mean correlation [Mann–Whitney *U*= 362, *Z*=2.63, *p=*.009, d_Cohen_=.666] and mean duration [Mann– Whitney *U*= 361, *Z*=2.60, *p=*.009, d_Cohen_=.658]; furthermore, marginally significant lower values were observed for the GEV [Mann–Whitney U= 358, Z=2.52, *p*=0.012, d_Cohen_=.636] and time coverage [Mann–Whitney *U*= 356, *Z*=2.46, *p=*.04, d_Cohen_= .621].

For the N250 (maps 5 and 6), group differences were found for both maps in both conditions. For direct gaze, ADHD patients showed higher values than controls in map 5 for mean correlation [Mann–Whitney *U*= 156, *Z*=-2.67, *p=*.008, d_Cohen_=.751], mean duration [Mann– Whitney *U*= 157, *Z*=-2.63, *p=*.008, d_Cohen_= .743] and time coverage [Mann–Whitney *U*= 150, *Z*=-2.80, *p=*.005, d_Cohen_= .799], as well as marginally significant lower values for the GEV [Mann–Whitney *U*= 161, *Z*=-2.54, *p*=0.011, d_Cohen_= .712].

For averted gaze, no differences were observed in the mean correlations between patients and controls (*p*>0.05). However, ADHD patients showed marginally significant higher values than controls in map 5 for GEV [Mann–Whitney *U*=185, *Z*=-2.06, *p=*.036, d_Cohen_= .533], mean duration [Mann–Whitney *U*= 173, *Z*=-2.40, *p=*.016, d_Cohen_= .621] and time coverage [Mann– Whitney *U*= 173, *Z*=-2.40, *p=*.016, d_Cohen_= .621]. Similarly, for averted gaze, marginally significant lower values in map 6 were found in ADHD than controls for GEV [Mann–Whitney *U*= 352, *Z*=2.00, *p=*.045, d_Cohen_= .591], mean duration [Mann–Whitney U= 361, Z=2.22, *p*=.026, d_Cohen_= .658], and time coverage [Mann–Whitney *U*= 361, *Z*=2.22, *p=*.026, d_Cohen_= .658]. Differences in mean correlations were not significant between patients and controls (*p*_*s*_ > 0.05).

For the P200 (maps 3 and 4), group differences were confirmed only for map 4 in the direct gaze condition. ADHD patients showed significantly lower map 4 values than controls for mean correlation [Mann–Whitney *U*= 362, *Z*=2.63, *p=*.009; d_Cohen_=.666], and mean duration [Mann– Whitney *U*= 361, *Z*=2.60, *p=*.009; d_Cohen_= .658]. No significant differences were observed in the GEV and time coverage between patients and controls (*p*>0.01). For the N250, group differences were found only for map 5 in the direct gaze condition. ADHD patients showed higher values than controls in map 5 for mean duration [Mann–Whitney *U*= 157, *Z*=-2.63, *p=*.008; d_Cohen_= .743] and time coverage [Mann–Whitney *U*= 150, *Z*=-2.80, *p=*.005; d_Cohen_= .799]. Differences in GEV were not significant between patients and controls (*p*_s_ > 0.01).

For averted gaze, no significant differences were observed in the GEV, mean correlations, mean duration, or time coverage between patients and controls (*p*>0.01).

### Effects of anxiety and correlations between clinical scores and microstates

To assess potential confounding effects of higher levels of anxiety in ADHD on eye-gaze processing (see (Rohner, 2002; Schmitz, Scheel, Rigon, Gross, & Blechert, 2012)), correlations were calculated between anxiety scores and microstates’ GEV. The GEV was selected as a representative microstate signature to minimize the number of multiple tests. Associations between variables were estimated using Pearson’s correlations (two-tailed), with bootstrapping interval estimations. For the direct gaze condition, STAI-trait scores were positively correlated with the GEV of map 6 (*p*=.018, *r* =-.307; confidence intervals [.054, -.79]; *r*^*2*^=.094). No other significant correlations were observed (all *p*_*s*_ > 0.5).

To the same end, an additional analysis was performed on extreme groups of trait-anxiety. First, a k-means cluster analysis was performed to categorize participants in three groups for levels of trait-anxiety. Subsequently, participants were distributed accordingly to each cluster: low (cluster center: 30.9), medium (cluster center: 40.52), and high (cluster center: 55.2). The two extreme groups (low vs. high) were then selected for subsequent analysis: 18 participants in the low group (14 controls and 1 ADHD), and 10 in the high group (9 ADHD and 1 control). Statistical analyses were re-performed on the maps and related parameters that were identified as different between healthy controls and ADHD: map 3 of the direct gaze condition (mean correlation, mean duration), and map 5 of the direct gaze condition (mean correlation, mean duration, and time coverage) (see Figure 3). Non-parametric statistics (Mann–Whitney *U* test) were used for analyses on these microstate parameters. *P-*values were corrected for multiple comparisons based on the number of estimated independent tests (see above).

**Figure 3.**
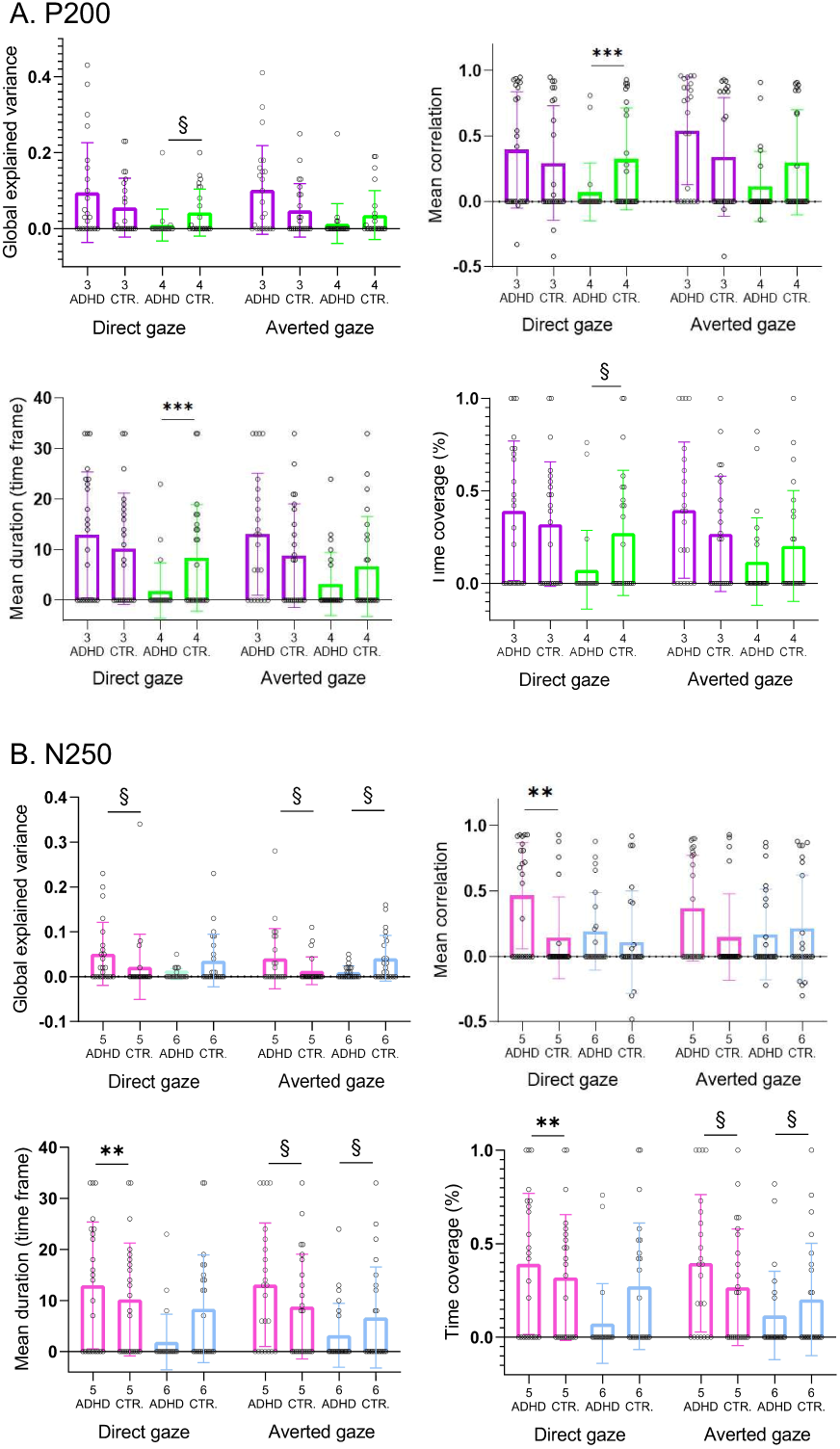
Microstate parameters of the P200 and N250. **C**omparison between groups for each microstate class on the global explained variance, mean correlation, mean duration, and time coverage. Significant group differences are marked by asterisks (§ *p*<.05 (marginally significant); \*\**p*<.01, \*\*\**p*<.005 (statistically significant)).

**Figure 4.**
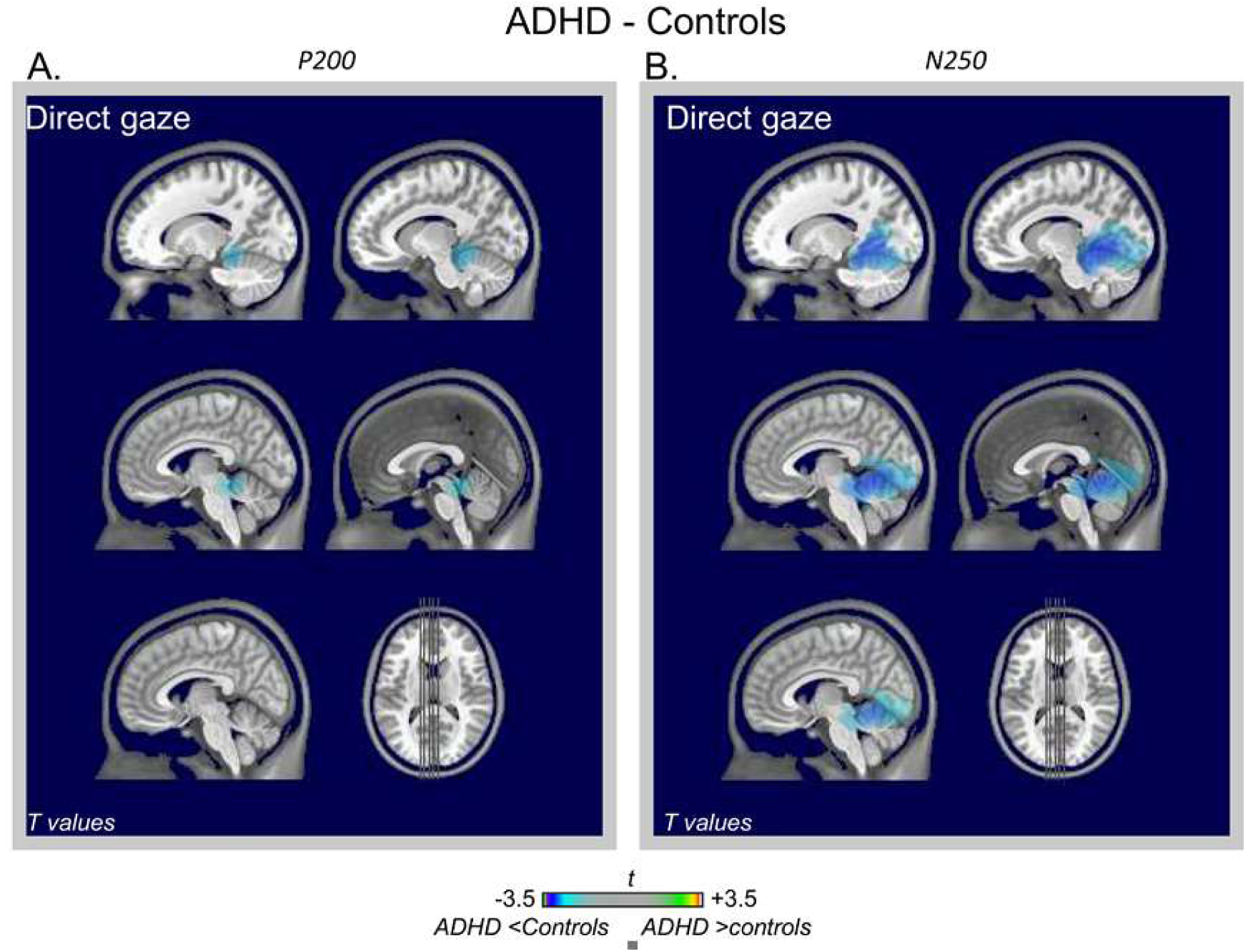
**A**. Differences in brain activation during attentional allocation to direct gaze between ADHD and control participants estimated for the P200 (microstates 3 and 4 time-windows), **B**. and the N250 (microstates 5 and 6 time windows). All activations showing statistically significant at *p* <.01.

For map 5 of the direct gaze condition, two effects were observed: individuals with low trait-anxiety showed marginally significantly lower values than individuals with high trait-anxiety for mean correlation [Mann–Whitney *U*= 138, *Z*=2.61, *p=*.021; d_Cohen_=.891], and significantly lower values than individuals with high trait-anxiety lower for time coverage [Mann–Whitney *U*= 134, *Z*=2.44, *p=*.009; d_Cohen_=.993]. No other significant differences were observed between the two groups (*p*>0.05).

To assess whether the reduced activity in P200 or N250 could predict the ADHD symptoms, mean durations of maps 4 and 6 (one of the strongest between-group effects, see Figure 3) were correlated with Inattention and Hyperactivity/Impulsivity scores. For the averted gaze condition, ADHD patients showed a significant negative correlation between mean duration of map 4 and Inattention/Impulsivity scores (*p*=.001, *r* =-.607; confidence intervals [-.846, .083]; *r*^2^=.368). No other significant correlations were observed (*p*_*s*_ >0.05).

### Brain imaging results

The analysis of microstates highlighted differences between groups at middle stage latencies. Time windows were defined by k-means clustering performed on the grand averages, as well as statistically significant differences between groups (see Figures 2 and 3). For the P200, group differences were confirmed only for the direct gaze condition. In the direct gaze condition, the randomization test showed differences between ADHD (time window: 168-236ms; microstate 3) and controls (time window: 168- 276 ms; i.e., microstate 4) in the left cerebellum and in the vermis (all *p*_*s*_ <.01). Reduced activations were found for ADHD patients compared with controls in all these brain regions [left cerebellum (*t=*-2.583, *p*=0.008; d_Cohen_=.762); vermis (*t*=-2.502, *p*=0.007; d_Cohen_=.738)].

For the N250, group differences were found only in the direct gaze condition. In this condition, by comparing ADHD (time window: 236-300 ms; i.e., microstate 5) with healthy controls (time window: 276-300 ms; i.e., microstate 6), the randomization test showed differences between groups in the left posterior cingulum, left calcarine, left and right lingual, left and right cerebellum and vermis (all *p*_*s*_ <.01). Reduced activations were observed for ADHD patients compared with controls in all these brain regions [left posterior cingulum (*t*= -2.446, *p*= 0.009; d_Cohen_=.721), left calcarine (*t=* -2.713, *p*=0.005; d_Cohen_=.8), left lingual (*t*= -3.045, *p*=0.003; d_Cohen_=.898), right lingual (*t*= -2.536, *p*=0.006; d_Cohen_=.748), left cerebellum (*t*= -2.974, *p*=0.002; d_Cohen_=.877), right cerebellum (*t*= -2.665, *p*=0.007; d_Cohen_=.786), vermis (*t*= -3.007, *p*=0.001; d_Cohen_= .887)].

In the averted gaze condition, the comparison between the ADHD sample (time window: 236-300 ms; i.e., microstates 5) and healthy controls (time window: 232-300 ms; i.e., microstate 6) resulted in no significant differences (all *p*_s_ >.01).

## Discussion

In this study, we have compared ERP markers of face and gaze processing in adults with ADHD and healthy controls. Microstate analyses allowed us to make hypotheses about dysfunctional temporal stages of eye-gaze processing in the adult ADHD brain. ADHD was found to be associated with distinct abnormalities on the attentional-sensitive P200 component, and on subsequent higher-order processing time windows (N250). Source imaging of the N250 and P200, for faces with direct gaze, revealed hypo-activations in brain regions that play a key role in social functioning in adults with ADHD. This study provides the first evidence of alternative neural strategies for eye-gaze cue decoding in adults with ADHD.

With respect to behavioral performance, *d’* was high in both groups, evidencing an adequate execution of the task. Replicating our previous findings (Berchio et al., 2017; Berchio et al., 2016), for all participants, faces with averted gaze were found to be more discriminable than faces with direct gaze. No group differences were found on indexes of face recognition. Similar results have been reported in previous studies (Borhani & Nejati, 2018; Ibáñez et al., 2011; Noordermeer et al., 2020). Furthermore, the absence of biases as assessed by *d’* and the *C* criterion means that we can be confident that our findings are not explained by distractibility/impulsivity in ADHD. RT data indicated that ADHD patients have a tendency to react faster to faces with direct gaze when compared to controls. Based on this marginally significant effect, we speculate that in adults with ADHD, direct gaze may affect perception, attention, and cognition. However, taken together, these findings indicate that the behavioral deficits so often observed in children with ADHD may be compensated by adopting alternative neural strategies in adults.

There is evidence that impulsivity and attention deficit are essentially modulated by primary intelligence in children with ADHD, and lower scores of intelligence are associated with higher levels of impulsivity (Buchmann, Gierow, Reis, & Haessler, 2011). In this study, scores collected with the two subscales of the WAIS, together with the K values, seem to exclude executive functioning deficits regarding the variables assessed in our task. Furthermore, since mental retardation was a criterion of exclusion, we can assume that a sample of patients with lower levels of intelligence would have shown a weaker performance due to impulsive responses.

From microstate analyses, we showed that ADHD adults and controls displayed similar P100 and N170 topographies for direct and averted gaze. The P100 is modulated by spatial attention linked to sensory-gating mechanisms of the visual cortex (Couperus, 2010; Gherri & Forster, 2015). This finding suggests that adults with ADHD present intact low-level processes related to visual orienting to faces. Previous researchers have suggested altered P100 responses during eye-gaze processing in children with ADHD (Groom et al., 2017; Tye et al., 2013). Despite the difficulty of comparing P100 sensory modulations using different tasks (Burra, Framorando, & Pegna, 2018), we may assume that through repeated exposure to social situations, adults with ADHD have developed better sensory strategies.

Impaired emotional facial processing is the most consistently reported form of social cognitive impairment in ADHD (Da Fonseca, Seguier, Santos, Poinso, & Deruelle, 2009; Miller, Hanford, Fassbender, Duke, & Schweitzer, 2011), and studies in adults with ADHD have indicated altered N170 responses to emotional faces (Ibáñez et al., 2011; Köchel et al., 2012). In our study, we found preserved N170 maps, likely due to the fact that our stimuli were neutral faces.

Topographical differences between groups were observed at the latency of the P200 and N250. For faces with direct gaze, map 4 was found to have reduced GEV, mean correlation, mean duration, and time coverage in the ADHD sample when compared to controls. One of the most prominent theories regarding the P200 is that it indexes networks associated with attentional selective processes (Ibanez et al., 2012; Wuttke & Schweinberger, 2019). The P200 appears also particularly suitable to assess attentional allocation to gaze direction and distinguish specific features of emotional dysregulation disorders (Berchio et al., 2017). Our results illustrate specific abnormalities in allocating attentional resources to a face with direct gaze in adults with ADHD. Putting results together, we may further assume that two competing attentional mechanisms for eye-gaze processing occur at this latency, probably reflecting different levels of attentional efficiency. The near absence of map 4 in the ADHD sample may indicate the predominance of an alternative attentional mechanism.

At later processing stages, adults with ADHD showed atypical ERP microstates in response to faces with both direct and averted gazes. At approximatively 250 ms, an enhanced posterior negative potential is considered to index higher-order cognitive processes, such as facial familiarity recognition (Pierce et al., 2011; Zheng, Mondloch, & Segalowitz, 2012). Previous ERP studies have reported impairments at middle latency stages when ADHD adults attend to target faces (Raz & Dan, 2015). Our results further support abnormalities in decoding eye-gaze in adults with ADHD.

Regarding the association between anxiety scores and microstates’ GEV, in the ADHD sample, we only observed a positive correlation between map 6 of direct gaze and traits of anxiety. Increasing arousal may be necessary to obtain a more in-depth evaluation of socially relevant signals. Nevertheless, this relation was not present in the prototypical map of the ADHD sample. Therefore, this finding may also be considered as an indication of an atypical affective reactivity to eye-contact in ADHD.

An additional analysis was also performed to further assess confounding effects of trait-anxiety. This analysis highlighted a significant effect, but again and importantly on the latency corresponding to the N250 for the direct gaze condition. This is not surprising since eye-gaze perception is modulated by anxiety, and the effects of arousal on direct gaze have been documented many times (Rohner, 2002; Schmitz et al., 2012). However, what is important to remark here is that both analyses exclude the impact of trait-anxiety on the P200. In the averted gaze condition, ADHD patients also showed a negative correlation between the mean duration of map 4 and inattention/impulsivity scores. This result indicates that higher levels of inattention may predict abnormal neural processing to eye shifts and may therefore also negatively impact social interactions.

We additionally performed source imaging on microstates corresponding to the latency of the P200. For faces with direct gaze, this analysis highlighted diminished activations in the left/midline cerebellum, in ADHD patients compared to controls. The cerebellum has long been known for its importance in motor learning and coordination (Mauk, Medina, Nores, & Ohyama, 2000; Miall, 1998; Roldan Gerschcovich, Cerquetti, Tenca, & Leiguarda, 2011), but growing evidence has also highlighted its crucial involvement in emotion recognition, theory of mind, and empathy (Adamaszek et al., 2015; Roldan Gerschcovich et al., 2011). More importantly, the cerebellum has emerged as one of the key dysfunctional brain nodes in ADHD (Bruchhage, Bucci, & Becker, 2018; Krain & Castellanos, 2006; Stoodley, 2014). Reduced cerebellar activity may lead to dysfunctionally integrating social inputs to an executive program. Moreover, our results may suggest that ADHD-related social impairments may be potentially explained by other concomitant factors, ‘such as an inability to engage in proper eye contact’. Future studies should investigate this causative link.

We additionally performed source imaging on microstates’ latencies corresponding to the N250. For faces with direct gaze, this analysis highlighted diminished activations in the bilateral cerebellum, visual regions, and the left posterior cingulate, in ADHD patients compared to controls. The posterior cingulate is involved in controlling internal balance between internal and external attention (Leech & Sharp, 2014), and together with the cerebellum, plays a key role in modulatory influence of cortical regions implicated in social cognition (Spreng & Andrews-Hanna, 2015; Van Overwalle, D’aes, & Mariën, 2015). Our finding of ADHD-related decrease activity in these posterior regions may suggest a locus of dysfunction in mediating internal aspects of social cognition in ADHD.

A limitation of this study is that the experimental design does not allow us to discriminate between emotions, and therefore conclusions may be drawn only to neutral eye-gaze processing. Another limitation is the small sample size, which limits the generalizability of the study’s findings and may have reduced the possibility of capturing subtle deficits, especially regarding behavioral effects. These data provide preliminary evidence in an under-studied population, but a larger sample size would have provided greater statistical power. Therefore, future studies should enroll a larger sample size to boost the clinical application of these results.

In conclusion, to our knowledge, this is the first investigation of the neural correlates of eye-gaze processing in adults with ADHD. The present findings indicate that this disorder in adulthood appears to be associated with brain impairments involving eye-gaze decoding. There has been sparse evidence for theory of mind deficits and reduced empathy in adults with ADHD. The impairments seen in middle latency stages may not be surprising since successfully processing social signals depends on our ability to accurately decode social meanings. The current study extends previous findings by demonstrating alternative neural mechanisms in processing socially relevant information, and it also indicates that brain deficits in attending to socially relevant information may persist across the lifespan. Evidence identified in this work can also be viewed as future directions for methodological research in the field of machine learning. The application of multivariate pattern analysis (MVPA) to ERPs has opened up new areas to understand how mental representations are processed in the brain (King & Dehaene, 2014). To the best of our knowledge, so far only one EEG resting-state study has tried to combine microstate analyses with MVPA (Baradits, Bitter, & Czobor, 2020). Although it is clear that more methodological research is needed, the combination of these two methodologies may represent an opportunity of enriching both approaches. Further work in this area is absolutely needed to improve the effectiveness of tools for ERP analysis in clinical research.

## Acknowledgements

The study was supported by the Swiss National Center of Competence in Research [“Synapsy: the Synaptic Basis of Mental Diseases” no.: 51NF40-158776; and grant no.: 32003B_156914]. The Cartool software is a freely available academic software that has been programmed by Denis Brunet (https://sites.google.com/site/cartoolcommunity/).

